# Epi-fluorescence Microscopy of Single Molecule DNA Denaturation *in situ*

**DOI:** 10.1101/368423

**Authors:** Pulkit Sharma

## Abstract

DNA can be denatured by two main methods which are: a) denaturation in solution *(invitro)* and b) denaturation on a slide surface *(in-situ)*. Additionally, DNA can also be denatured in gels with urea. The method to be used depends on various factors such as the application, the source of the DNA, the length, and the techniques available to confirm the extent of denaturation. Verification of the extent of denaturation is important because of the following factors: 1) increases the chances of hybridization (especially for short probes), 2) prevents the loss of expensive probes (if the target site is not denatured then, the probes will not hybridize and will only cause a high a background), 3) a higher degree of denaturation allows for more probes to be used and therefore, more information can be derived after hybridization, and 4) essential to maximize due to extremely short probe length. It is important to ensure that DNA morphology is preserved after denaturation in order for the probes to hybridise and also for ensuring proper statistical analysis for high throughput applications. In this work, various experimental conditions for in situ denaturation of single molecule DNA is presented.

**Significance Statement:** The significance of this work is that it emphasizes on the importance of denaturation of target genomic DNA in DNA fibre FISH (fluorescence in situ hybridisation) experiments. If the quality of the target DNA is poor after denaturation or the target DNA is not properly denatured, then it will be very difficult or impossible to hybridize the probe DNA during FISH experiments. This will affect the final results for DNA FISH. Additionally, it is the first time that single DNA combed molecules have been shown to be denatured in situ. Most of the past work has been on gels only. Thus the work is both unique and significant.

## Introduction

Several studies have been done on DNA denaturation in different conditions by many research groups. Each study has its own purpose for denaturing DNA. In this study, the aim is to find out the best denaturation conditions for lambda DNA *in vitro* and *in situ* for future hybridization experiments. The extent of denaturation along a single DNA molecule needs to be maximized for maximum hybridization of probes. DNA dentures following mild treatment such as low pH, high temperature and low salt concentration) (Thomas, 1993). DNA melting is the separation of the two strands of the double helix by heating, or changing other parameters such as salt concentration or hydrostatic pressure [20], [21] (see Figure 1). It is possible to monitor DNA melting since the breaking of the hydrogen bonds between base pairs is accompanied by a very large increase of the absorbance of ultraviolet light at nearly 260 nm. Additionally, denaturation occurs in multiple steps and it is highly sensitive to the given sequence order (Hernandez-Lemus, 2012). Formamide is a good denaturant in low temperature conditions (Blake, 1996). According to Singh et al., HCl denaturation discriminates markedly against GC rich DNA. Chromosome morphology is best preserved after HCl and heat denaturation. (Singh 1977). After denaturation, changes occur in scattered light flux and optical density. (Dubey, 2005). The denaturation rate of double stranded DNA decreases exponentially as a function of length below the denaturation temperature (Erp, 2012). DNA’s double helix stability and stiffness are inversely proportional to the urea concentration. (Zhu, 2016). Formamide is often used as a denaturant since it influences the DNA melting (Rauch 2016). Krasna et al have shown that the continuous addition of acid or alkali to maintain a DNA solution at*p*H 7.0 results in the irreversible denaturation of DNA (Krasna, 1969). If the DNA is uniformly stained with a dye that unbinds when the DNA melts, local fluorescence of the melted region will decrease (Reisner, 2010). We have also shown this phenomenon in this paper in the results section. Additionally, the DNA double helix can be split into separate strands as a result of interaction between light and an intercalated dye. (Bernas 2006). We have also observed this as photocleavage of the DNA strands. While comparing DNA and RNA metling, the enthalpy change associated with melting of free rRNA [mean value, 30 J (g RNA)-l] is less than that for DNA [62 J (g DNA)-’ 1, due to differences in the base pairs (Seymour, 1991). Freezing has little effect on the degree of denaturation (Galyuk 2012).

Like Zen et al have reported, we have also observed the local opening of DNA bubbles in selective regions without breaking the whole DNA (partial denaturation)(Zen 2015). For thermal denaturation, DNA strands are separated at a high temperature (Iwasaki, 2004). Melameli et al, have studied the molecular mechanisms that are responsible for the differences in DNA sensitivity to denaturation in condensed versus diffuse chromatin (Melameli, 1987). Thus, many groups have studied the phenomenon of DNA denaturation in detail under various conditions.

## The following materials and methods sections describe various denaturation conditions and their associated results

### Section 1: Optimisation of denaturation of DNA in solution

#### Aim

To find out the best denaturation conditions for lambda DNA in solution for future hybridization experiments.

#### Background

The denatured regions of combed lambda DNA will eventually be targeted by complimentary oligonucleotide probes. Therefore, it is essential to optimize the denaturation conditions so that the sequence of interest is accessible to probing. In this section, a series of denaturation conditions in solution have been tried in order to improve upon previous experiments. The denatured areas can be confirmed by a number of means such as Acridine Orange staining (distinguishes between ssDNA (red fluorescence and dsDNA (green fluorescence)), AFM, ssb, and of course, hybridization itself.

#### Materials

Tris-EDTA (TE) pH 8.0, 0.1M NaOH, 70% formamide/2 X SSC, MilliQ water, X-tra slides, Matsunami coverslips, Sybr gold (1:500 dilution), lambda DNA, antifade (300ul 50% glycerol, 100ul AMP (AMP = 2-Amino-2-Methyl-1,3-Propandiol) 100ul 25mM Tris/EDTA/MgCl_2_, 100ul 20% 2-Mercapto-ethanol/TrisEDTA), 1mM EDTA, and 10mM Hepes/1mm EDTA.

#### Methods

A series of denaturation reactions were setup as follows:

**Table 1:** Buffers for Denaturation in Solution.

| **Tube:** | **Buffer (1ml)** |
| --- | --- |
| A | Tris-EDTA pH 8.0 |
| B | 70% formamide/2 X SSC |
| C | 0.1M NaOH |

All tubes contain 2ul lambda DNA (500ug/ml) in addition to their respective buffers.

*<u>Tube A:</u>* All the samples above were heated in solution for 5 minutes at 55°C and snap cooled.

**Note:***In sections A and B combing was done in the standard way as follows*:

15ul of each solution were combed between 2 Matsunami coverlslips and 10ul of antifade and 1ul of Sybr gold (1:500 in MilliQ water) was placed X-tra slides and the respective coverslips were placed on top for microscopy.

#### Section 1 Results

(The sections below corresp0ond to the sections in the methods above and the respective buffers are indicated in brackets for each pair of images):

**Figure 1.1:**
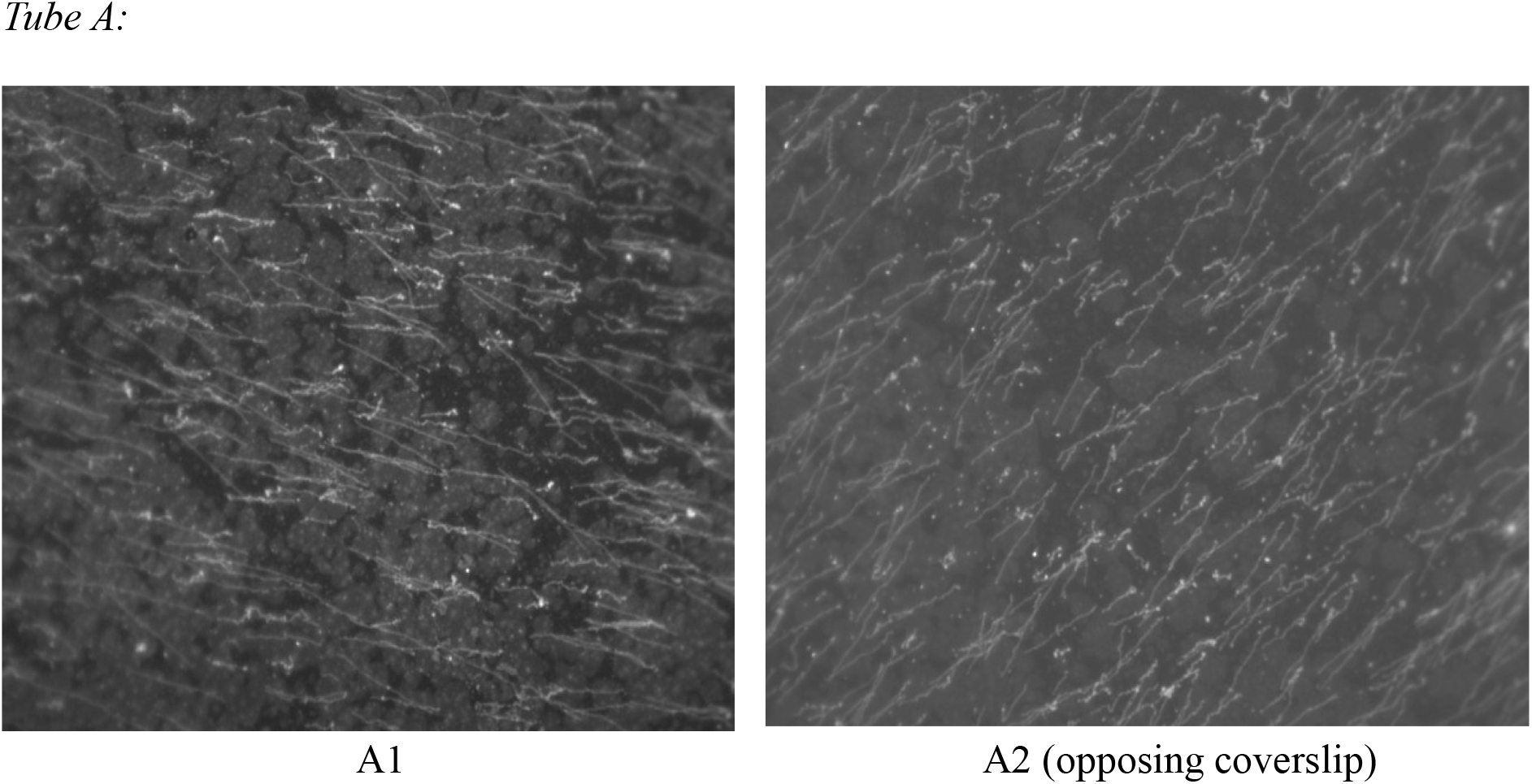
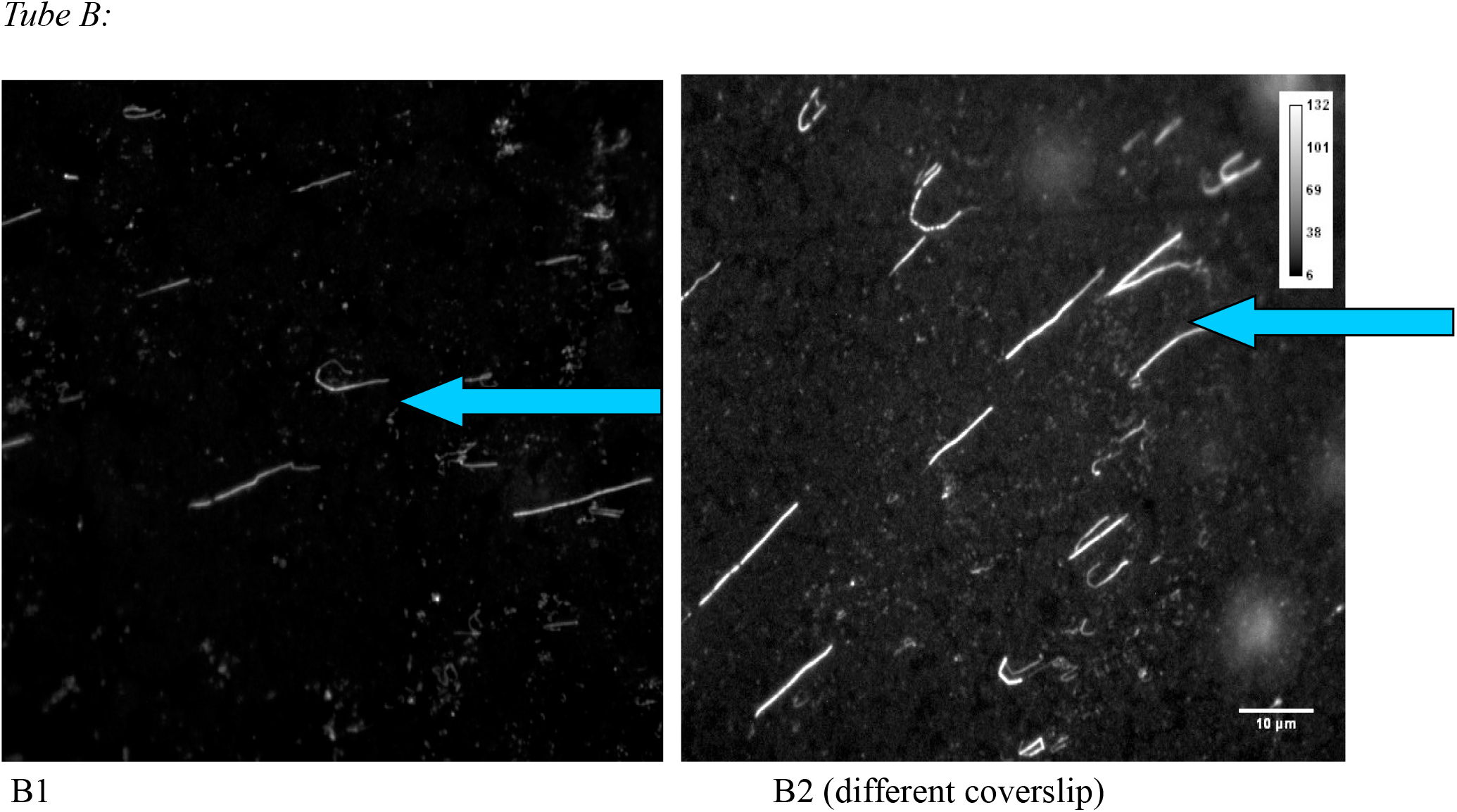
Denaturation at 55°C/5minutes. Denaturation is more evident in both B1 and B2 than in the previous coverlsips. “Partial” denaturation can be seen (arrows). Since the denaturation of lambda DNA seems to be the most successful in image D2 above among all the others, a surface plot and acalibration bars is provided to give an idea of the degree of saturation of pixels which is directly proportional to the fluorescence intensity. Most of the DNA is destroyed in B1 (70% formamide/2XSSC). Arrow in surface plot below corresponds to the area marked by an arrow in the image D2 above. Arrow in B1 also shows partial denaturation.

**Figure 1.2:**
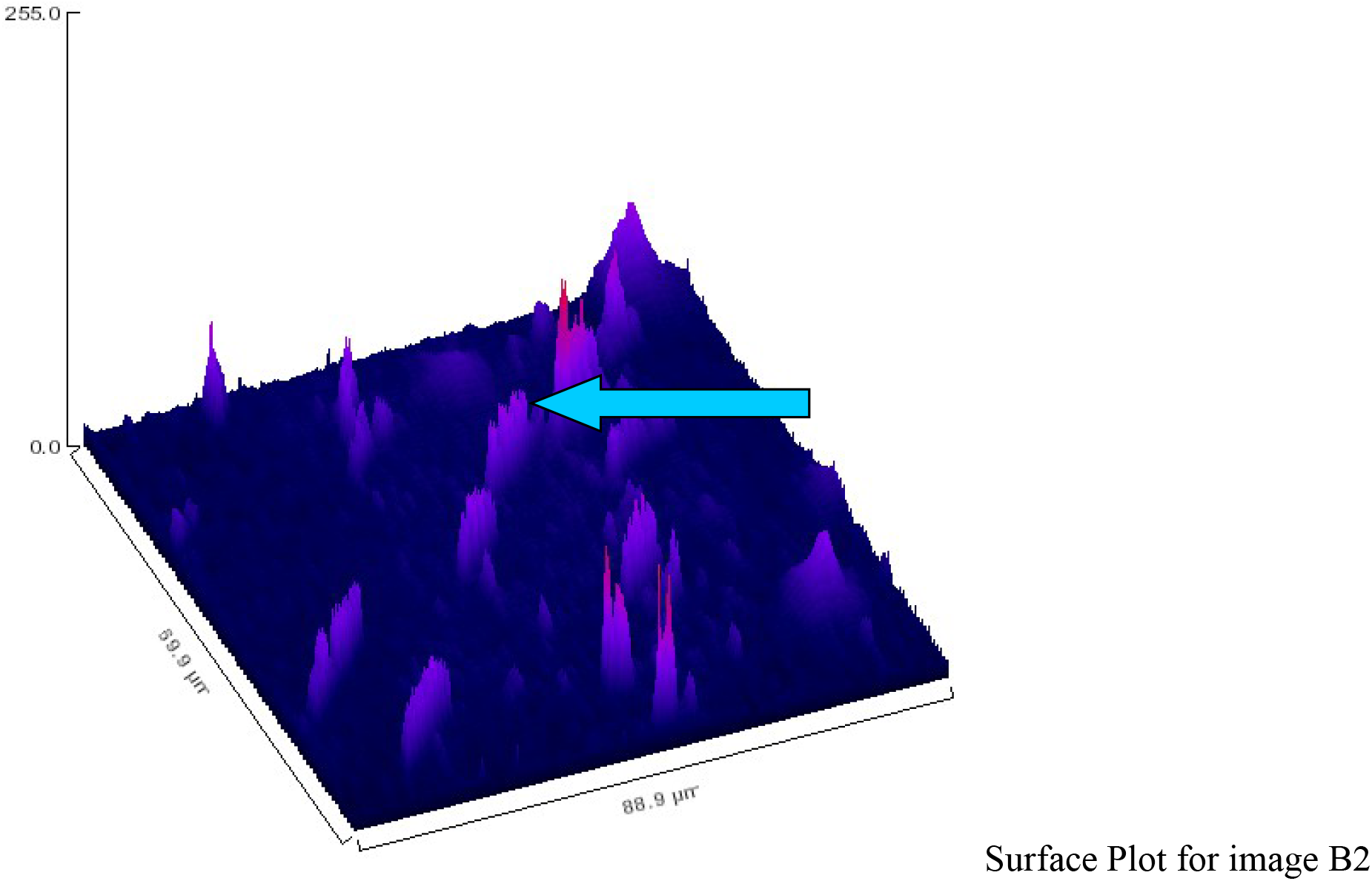

**Figure 1.3:**
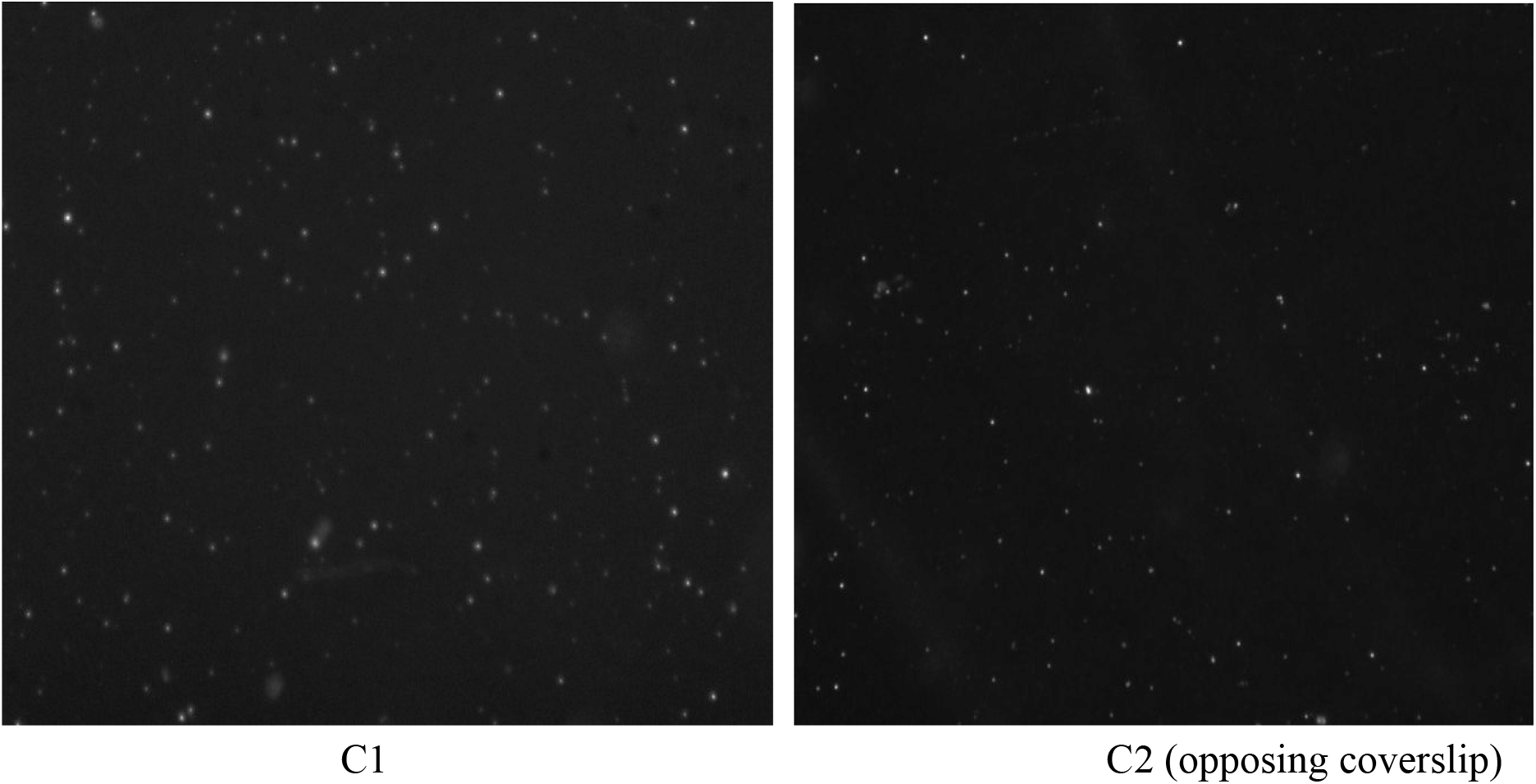
Tube C. All DNA destroyed. NaOH concentration may have been too high.(0.1M NaOH)

#### Discussion

The slides and coverslips were not cleaned and this could have led to nuclease action and contamination which affected the results. Both coverslips and slides should be thoroughly cleaned before experimentation. The sybr gold staining was not homogenous in some areas and could be confused with denaturation. However, I think that the denaturation is more evident when parts of the same molecule are lighter of darker in contrast to different fields of view on the same coverslips. A series dilution of ssb could confirm the identifation of ssb on a slide/coverslip. The controls showed bad combing and this could be due to the buffer being contaminated. A lower concentration of NaOH should be tried for a shorter period of time. It is noteworthy, that the combing and denaturation vary on the same coverslip (as seen from the different fields of view on the same coverslip) and on the opposing coverslip even though the conditions are the same. This suggests that the DNA needs to be spread and fixed on the coverslip surface more evenly and the heat transferred over a given area is affected by this non-homogenous spreading. There may be more fluid trapped in some areas than others causing uneven denaturation.

#### Conclusion

It is not evident from the results above if ssDNA molecules can distinguished from dsDNA molecules after denaturation. However, the lighter parts of the same DNA molecule after denaturation suggests that we can identify denatured regions. More optimization is required. MilliQ water and 70% formamide/2XSSC have shown the best overall denaturation results, however, denaturation with these still needs to be improved.

### Section 2: In-Situ Optimisation of Denaturation of combed DNA with a Single Buffer (*in-situ* Denaturation)

#### Aim

This experiment has the following aims: a) to check the extent of denaturation and to compare different conditions for denaturation of fast-combed DNA in order to optimize the conditions for fast combing, denaturation, and a low background. and, b) to check the quality and quantity of combed DNA after various treatments. The extent of denaturation can be checked by measuring the fluorescence intensity of the combed DNA before and after denaturation. It is expected that the fluorescence intensity would be roughly half for denatured DNA (single stranded DNA/ssDNA) than that of non-denatured DNA (double stranded or dsDNA). However, since renaturation can occur, the hybridization of probes along the length of the DNA molecules is the preferred method of ensuring that denaturation has been optimised.

#### Reasoning

This experiment is important because it will establish the conditions essential for hybridisation of oligo probes to the target DNA molecules. It has been noted on several occassions that the initial problem of non-hybridisation lies within the various treatments the target DNA has gone through before adding the probes. It is essential to ensure that the target DNA is well combed out, properly denatured, and occurs in sufficient numbers. Often, target DNA is found to be broken (photocleaved or photobleached), poor quality (initially), washed off completely, or damaged after various treatments. This renders it unfit for probing. Thus, proper measures should be taken to ensure that probing is not begun until the quality and quantity of DNA is checked. This will save time and reagents. The extent of denaturation needs to be optimised while preserving the quality of the individual DNA molecules. Additionally, too many molecules, lead to large masses of DNA bundled together, concatermerization, or improper combing. Thus, the right density per unit area of the slide is also important.

#### Materials

X-tra slides, Matsunami coverslips, lambda DNA, antifade with sybrgold (1:200 dilution) per slide/coverslip, and 10mM Hepes/1mM EDTA, 1 X PBS /1%tween-20, 70% formamide.**Methods:** The combing solution was made as follows:

979ul 10mMHepes/1mM EDTA, 1ul lambda DNA (500ug/ml), and 20ul of 1:200 sybergold in water and 20ul of this solution were placed between two coverslips which were then pulled apart.

Denaturation in all cases was done for 2 minutes (50ul of 70% formamide/70°C on a heat block per coverslip/slide pair). In the following (sections A to C) fast-combing conditions were tested in which two matsunami coverslips each were used with X-tra slides as support for microscopy:

A) 2 X-tra slides (A and B) were used as support with antifade added on the slides after fast combing on matsunami coverslips *before* denaturation. After imaging, the coverslips were removed from the slides by dipping in 1 X PBS/1%tween (20 minutes) and then 50ul of 70% formamide was placed on the same slide followed by a fast combed coverslip for two minutes at 70°C. After removing the formamide, 10ul of antifade was added to the slide and the same cover slip was placed on top. More images were then taken.
B) 20ul of combing solution was placed between 2 cover slips (C and D) and fast combing was done after denaturation on the coverslips (i.e., coverslips were pulled as in fast combing *after* denaturation). Antifade was added after denaturation. Two X-tra slides were used as support for imaging.

#### Results

**Section 2:**
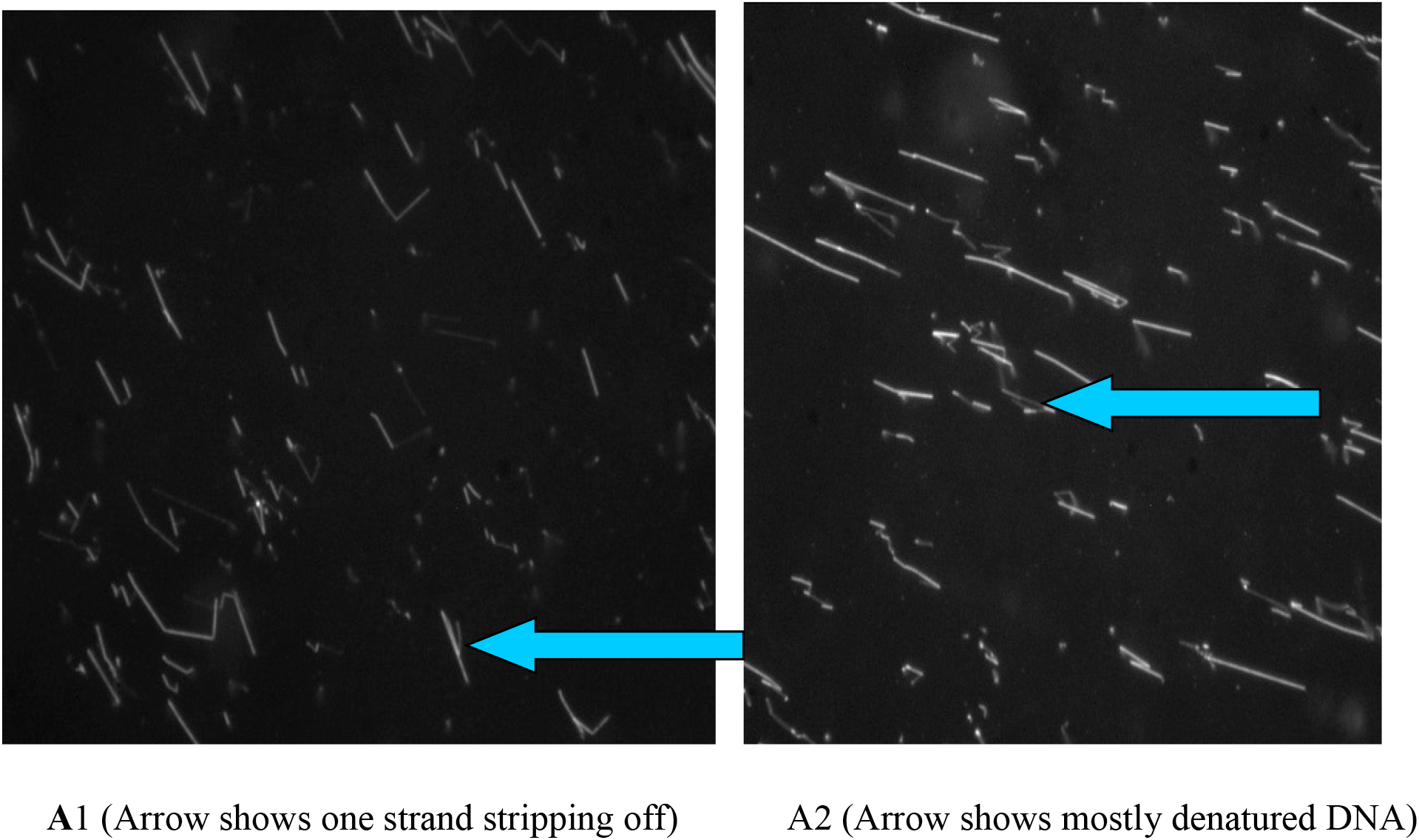
A1 shows denaturation on coverslip with antifade added afterwards whereas, A2 is a different field of view and shows some denatured molecules with a higher fluorescence intensity and low background.

**Section C: Figure 2.2:**
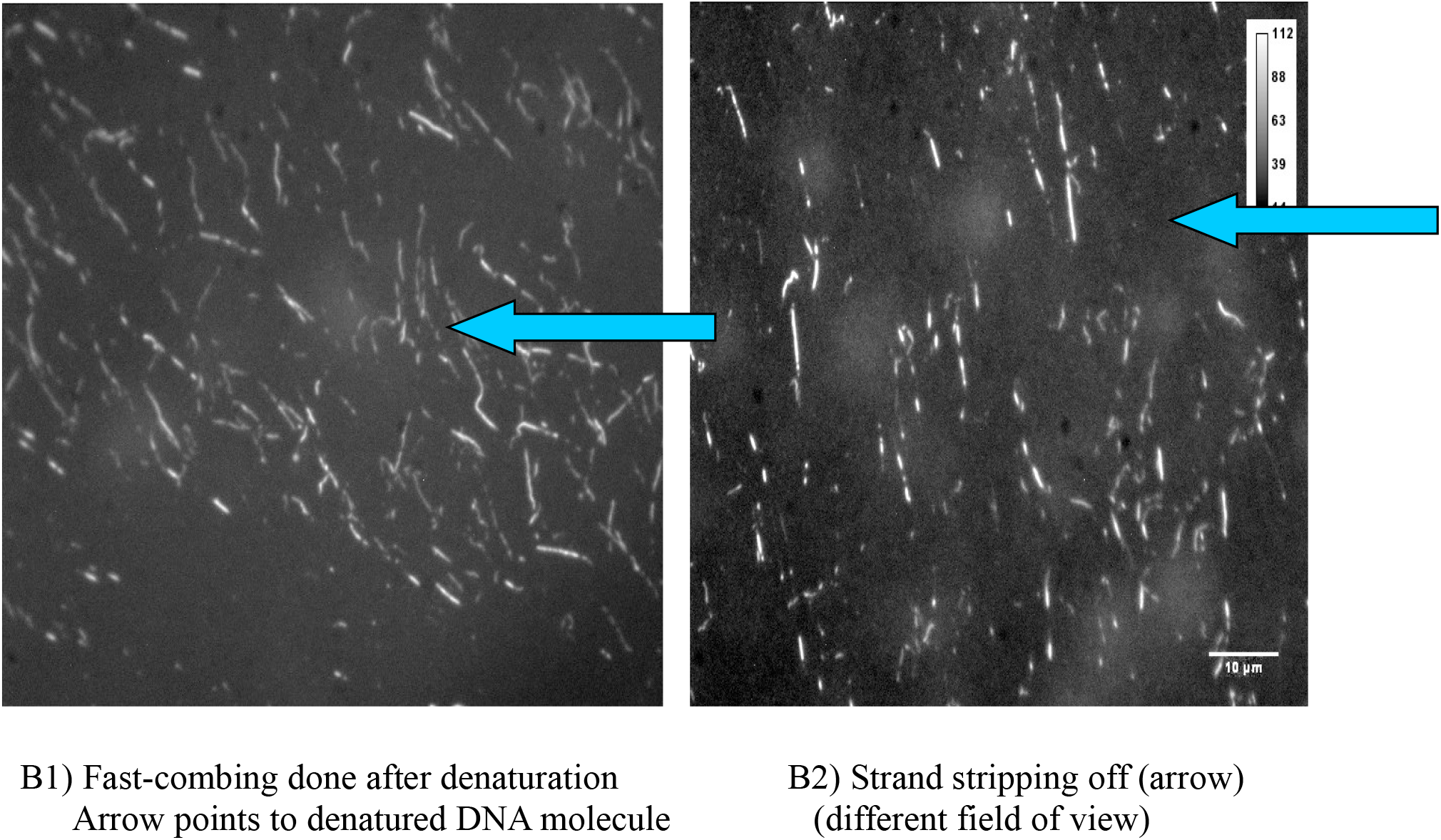

**Figure 2.3:**
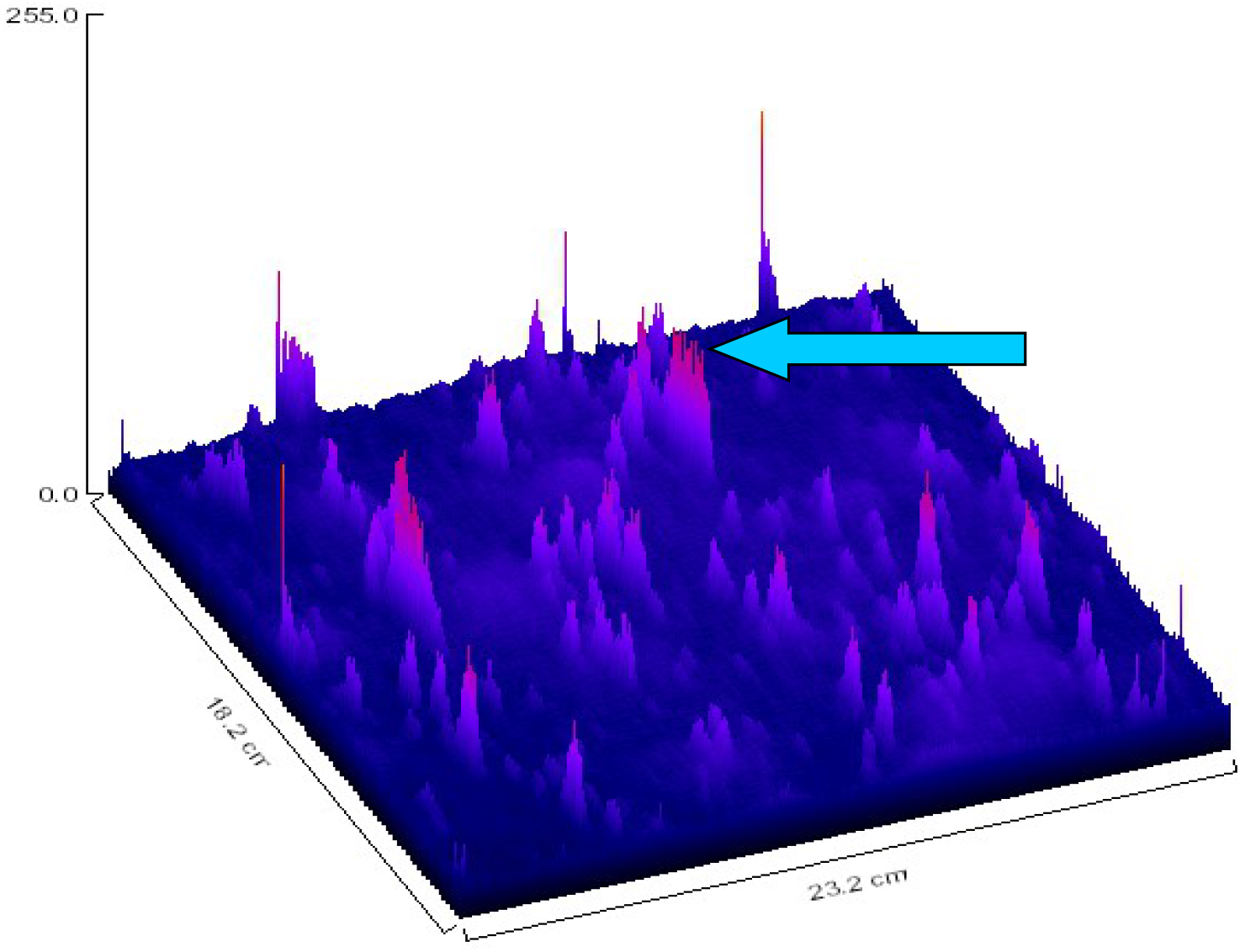
Surface plot for B2 (B1 and B2 show the best denaturation among all others coverslips)Arrow points to peak which represents DNA strand stripping off in image B2 above.

#### Discussion

In A and B, many stretched DNA molecules can be seen. They are the controls. Many molecules are wrinkled and different areas of the slide are not homogenous. A further 10X dilution could make the slide more homogenous. After denaturation (A and B), most of the DNA has been washed off and very few molecules are seen. Some of them may be denatured. Denaturation could be checked with enzymes or oligonucleotides. In C there is less DNA than D but in both cases there is not much denaturation and the combing is not homogenous. There are some faded areas. C and D have more DNA than A and B. Therefore, the antifade has apparaently allowed less DNA to stick (as in A and B) to the slide surface but it has hindered denaturation (as in C and D). E1 and E2 show the best denaturation. The fluorescence intensity of denatured versus non-denatured parts should be compared. The fluorescence intensity of the denatured part should be about half that of a non-denatured part of a stretched DNA molecule. The differerences in denaturation between different coverslips could be due the variations in the surfaces of the supporting slides or it could be an indication of non-homogenous denaturation. In other cases, the differences in denaturation within a coverslip pair can also be due to the position of a particular coverslip (ie, the one which is in contact with a heat block or the lower coverslip, will show a high degree of denaturation than the the one placed above it.

#### Conclusion

From A and B we can say that the matsunami cover slips don’t hold the DNA too well after denaturation. However, the use of antifade seems to help the attachment. Denaturation on coverslips seems to be better than on slides. This could be due to more heat absorbed by the slides. There is a need for more homogeneity in both combing and denaturation. The background is not good on the X-tra slides. From E1 and E2, it is evident that the ESCO coverslips with suitable denaturation conditions (70°C/70% formamide/2minutes) help maximally with denaturation.

### Section 3: Differentiation of ssDNA and dsDNA by Acridine Orange Staining

#### Backround

In order to differentiate between ssDNA and dsDNA after denaturation the dye acridine orange (A.O.) was used. The exitation and emission wavelengths are 500nm and 604nm respectively for ssDNA and 500nm and 514 resptectively for dsDNA.

Methods: The DNA combing solutions was made as follows:

<u>Solution A:</u>
5ul lambda DNA
20ul TE buffer pH 8.0
20ul 20mM MgCl2
10ul 2-amino-3-propandiol
5ul glycerol (50% in MilliQ water)
5ul triton-X 100 (0.1% in MilliQwater)
100ul MilliQ water
50ul 2-Mercaptoethanol (20% in TE buffer pH 8.0) + 10ul glycerol (50% in MilliQ water).

#### Denaturation

15ul of Solution A was placed on 1% APTES coated slides and a coverslip was placed on top and then this unit was denatured by dipping in 0.1M NaOH at room temperature (at a highly alkaline pH of 12.3) for 5 minutes. This was followed by a wash in 1 X PBS for 3 minutes and drying was done at 37C for 5 minutes in Techne. Then acridine orange (30ul of acridine orange in citric acid/pH 2.6 + 20ul of 2-amino-3-propandiol + 10ul of 50% glycerol in MilliQ water) was added and a new coverslip was placed on top. Then microscopy was done after 5 minutes incubation at room temperature.

#### Results

**Figure 3:**
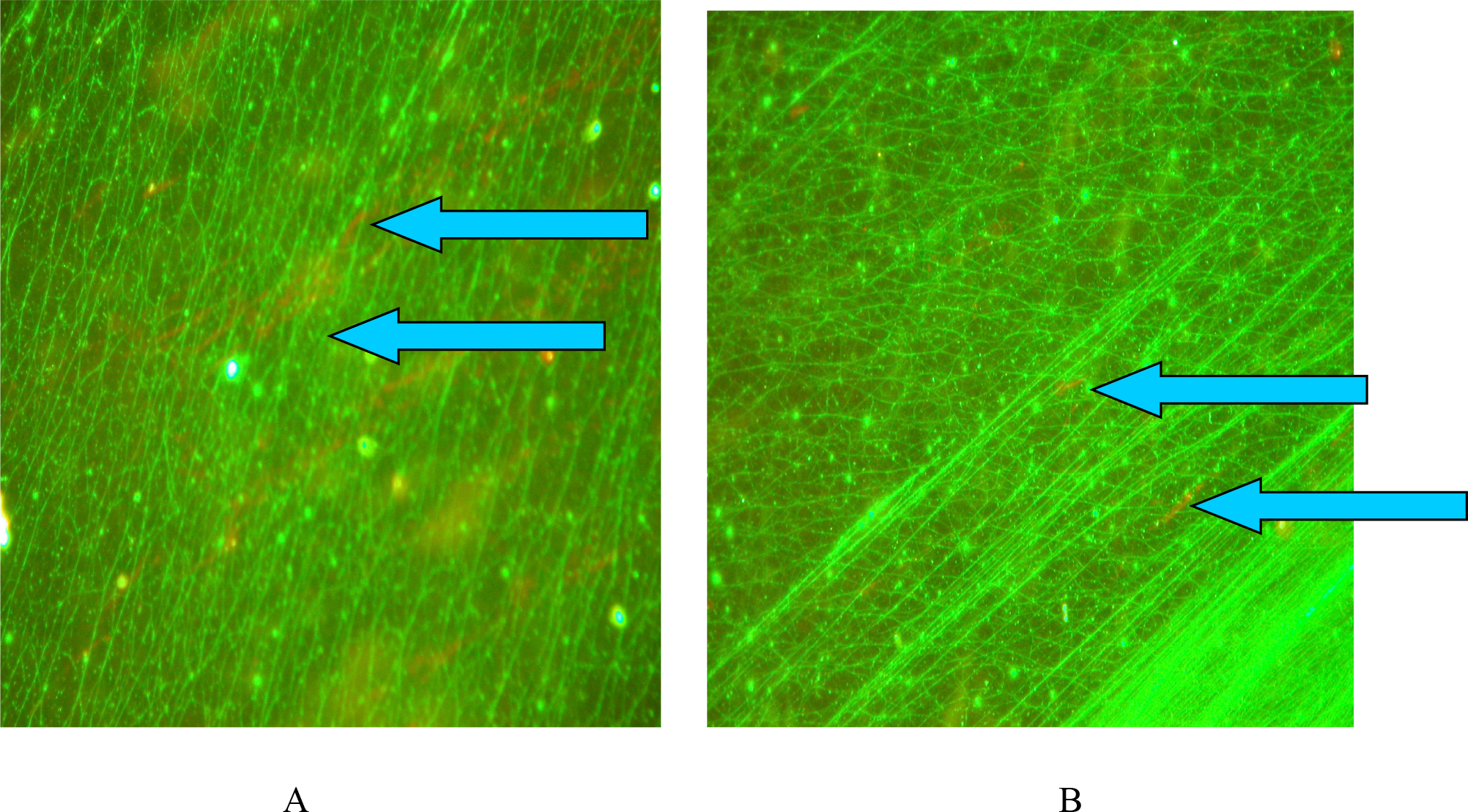
Solution A: Figures A and B show several green DNA fibres (dsDNA with a some denatured areas in red (arrows)

#### Discussion

These results show that the NaOH treatment is more effective than the heat block treatment. Additionally, The glycerol in solution Bmay have helped to preserve the morphology of the stretched DNA fibres. In some fields of view the red fluorescence was seen being rapidly replaced by green. The DNA concentration should be reduced (at least 10 times). Similar results have been reported by another group. There findings are as follows: “We had similar observation that red fluorescence is rapidly fading upon illumination and may be replaced by green fluorescence. I do not have an explanation of this phenomenon. It appears that red fluorescence is in fact the emission representing inter-system crossing (triplet excitation), perhaps a result of nuclei acid condensation (transition to solid state; the same happens to AO in solution upon freezing. There may be a problem in using AO on isolated DNA. The dye interacts with ss DNA by inducing its condensation – transition to solid state. The ‘dots’ that you see may be such condensates. Furthermore, at higher concentration (>20 uM in O.15 M NaCl) it may also induce denaturation of ds DNA, which then also condenses. It is a very ‘tricky’ dye that works quite well in aqeous solutions and in situ, but only at exactly right concentration and ionic strength, in equilibrium with DNA, when indeed it may differentiate between ss and ds DNA.” (email communication: Zbigniew Darzynkiewicz, et al, Brander Cancer Research Institute New York Medical College). They also suggested that Acridine orange should be purified by a membrane filter before use and a special laser/optical setup is required for proper imaging.

#### Final Discussion

The EMSA staining kit (for the gel shift assay with ssb: Molecular probes), ATTO-dino series of dyes (Atto-tec GmBH, Germany), and AFM analysis of combed and denatured DNA are some of the means to finally confirm denaturation. The EMSA kit will enable the simulataneous comparison of ssb protein-bound denatured DNA and unbound dsDNA. Fluorescence intentsity measurements over a statistically significant range of denatured and non-denatured combed DNA and its proper calibration with the appropriate software and microscopy equipment are needed in order to confirm if the pixel saturation values provided by a particular software are reliable. In my experiencec, the addition of glycerol based antifade solutions. Therefore, it may be advisable to use a solution of 1:200 Sybr gold dye in MilliQ water.

